# Global effects of *ade8* deletion on budding yeast metabolism

**DOI:** 10.1101/2021.03.15.435510

**Authors:** Agnese Kokina, Kristel Tanilas, Zane Ozolina, Karlis Pleiko, Karlis Svirksts, Ilze Vamza, Janis Liepins

## Abstract

Purine auxotrophy is a typical marker for many laboratory yeast strains. Supplementation of additional purine source (like adenine) is necessary to cultivate these strains. If not supplied in adequate amounts, purine starvation sets in. We tested purine starvation effects in budding yeast *Saccharomyces cerevisiae ade8* knockout. We explored effects brought by purine starvation in cellular, central carbon metabolism and in the global transcriptome level.

We observed that cells cultivated in purine depleted media became significantly more tolerant to severe thermal, oxidative and desiccation stresses when compared to the cells cultivated in media with all necessary supplements. When starved for purine, cells stop their cell cycle in G1 or G0 state; intracellular concentration of ATP, ADP and AMP decreases, but adenylate charge remains stable. Intracellular RNA concentration decreases and massive downregulation of ribosomal RNA occurs.

We think that purine auxotrophic starvation in a way mimics “natural” nitrogen or carbon starvations and therefore initiates elements of a transcriptional program typical for stationary phase cells (cell cycle arrest, increased stress resistance). Therefore our results demonstrate that organised metabolic response is initiated not only via “natural starvations”, but also when starving for metabolic intermediates, like purines.

## Introduction

Budding yeast *Saccharomyces cerevisiae* is an unicellular fungus that has evolved for fast growth in nutrient abundance (Duan et al., 2018). Deprivation of certain nutrients for yeast could be interpreted as a prelude of potentially more serious stressors to come. Lack of nutrients drives cells to enter quiescence state, where they do not proliferate, but become more stress resistant (Daignan-Fornier & Sagot, 2011, Gray et al., 2004). In case of carbon, nitrogen or phosphorus depletion, yeast cells stop cell cycle and activate specific gene expression patterns, thus forming “general stress resistance phenotype” (Gasch et al. 2000, Brauer et al. 2008, Klosinska et al. 2011). Lack of carbon, nitrogen or phosphorus initiates so-called “natural starvations” as yeast cells can experience those in “the wild”. In large, genome scale phenotyping screens, many mutations have been identified to increase budding yeast fitness when starved for carbon or nitrogen (Galardini et al. 2019).

*S. cerevisiae* is represented by a number of isolates or strains: many of them are found “in wild” but some of them are model strains or “laboratory strains”. S288c, CEN.PK and W303 are one of most popular budding yeast laboratory strains (Wang, et al., 2012). Among them, CEN.PK series have been specifically designed by K.D. Entian and P. Kotter and these derivatives are often used as models for fundamental physiology or industrial processes research (Entian & Kotter, 2007). Some characteristics of the CEN.PK strain series set it apart from others laboratory strains. In contrast to S288C, which cannot consume maltose anaerobically, CEN.PK113-7D has a functional MAL locus. In contrast to S288c and W303, CEN.PK strains are biotin prototrophs (Nijkamp et al. 2012). Wild strains are usually prototrophic for all nucleotides and amino acids, but many “laboratory strains’’ are auxotrophic (Pronk 2002). To use the CEN.PK series in gene engineering applications, specific auxotrophic markers are introduced. CEN.PK2-1D has *leu2-3/112, ura3-52, trp1-289* and *his3* mutations leading to leucine, uracil, tryptophan and histidine auxotrophies (Entian & Kotter 1998, Entian & Kotter, 2007).

When setting up cultivation experiments, often it is important to reach an appropriate growth rate, biomass yield and avoid unexpected phenotypic side effects. Therefore, proper concentrations for all auxotrophic nutrients in the media should be ensured (Pronk 2002). Alternatively - sudden depletion of auxotrophic nutrient might initiate specific, auxotrophic starvation and thus induce a number of phenotypic effects in yeast cells (Puig-Castellvi et al., 2016, Petti et al., 2011, Boer et al., 2008, Boer et al., 2010). Due to introduced auxotrophic markers in laboratory strains, specific “synthetic starvation” might set in when a specific auxotrophic nutrient is not supplied or is exhausted. Gomes et al. 2007 shows that limitation of essential auxotrophic amino acids decreases yeast final biomass yield and stress resistance. In the case of uracil or leucine starvation cells fail to enter a quiescence state and mostly die in an exponential growth phase (Boer et al., 2008, Brauer et al., 2008). On another hand, methionine starvation has shown signs similar to natural starvation response (Petti et al. 2011, Sutter et al. 2013), that is explained by methionine being a source of sulphur. This shows that not all auxotrophies are the same and can leave dramatically different effects on yeast metabolism. Also, strain genetic background can affect phenotypic response to environmental changes (Gomes et al 2007, González et al. 2007, Young and Court 2008, Alam et al., 2016, Galardini et al. 2019). Therefore, to continue exploiting model yeast strains in fundamental or applied research, detailed knowledge on their physiology in every possible environmental or laboratory setting is invaluable.

Although adenine auxotrophic strains are widely used, their phenotypic response to purine starvation is not studied in detail. Purines are ubiquitous molecules in the cell and form DNA, RNA (adenine and guanine nucleotides), as well as cofactors (NAD, FAD). Purine synthesis is highly conserved among eukaryotes. In budding yeast it consists of a linear chain of 9 sequential reactions coded by ADE1/2/4/5,7/6/8/12/16,17 genes. This pathway produces inosine monophosphate (IMP) which is a branching point to adenine, guanine and hypoxanthine. *ADE8* codes for phosphoribosylformylglycinamidine synthase which is the third enzyme in the chain. Currently no specific regulation activity of *ADE8* on IMP production is known (Rebora, et al., 2005). If *ade8* mutant are placed in the adenine deficient media, it cannot produce IMP, thus neither adenine or guanine is produced and purine auxotrophic starvation sets in.

Until now, we have demonstrated some purine starvation effects on *ade2* strain in the W303 strain background. In the case of purine starvation, W303 *ade2* yeast became desiccation tolerant, budding index decreased and trehalose content increased (Kokina et al., 2014).

In this article we report effects caused by purine starvation in purine synthesis in *ade8* knockout in CEN.PK2-1D strain. We analyzed various aspects of cell phenotype: cell growth, cell cycle state, changes in central carbon metabolism, cell ATP content, sublethal stress resistance and genome wide transcriptomic response.

Our results imply that purine auxotrophic starvation initiates formation of stress resistance phenotype and switches to fermentative growth. The transcription pattern of purine starvation resembles 48 h stationary phase cells that have ceased to multiply due to “natural starvation”.

## Materials and methods

### Strains and cultivation conditions

Wild type strain CEN.PK2-1D MATalpha *his3Δ1; leu2-3_112; ura3-52; trp1-289; MAL2-8c; SUC2* - gift from Phd. Peter Richard, VTT Biotechnology, Finland.

*ade8* CEN.PK2-1D *ade8Δ0. ade8* knockout was introduced by *ura3-URA3* 5-FOA toxicity knockout technique, using *ade8* knockout construct from Sadowski et al. 2008.

Cultures were maintained on YPD agar and kept in 4°C. Fresh YPD agar plates were regularly reinoculated from stock cultures kept in −80°C.

Strains were cultivated in SD media (Saldanha et al. 2004) with added 80 mg/L tryptophan, 100 mg/L uracil, 480 mg/L leucine, 100 mg/L histidine and 100 mg/L adenine as suggested by Pronk 2002.

To ensure that yeast culture is in exponential growth phase, we reinoculated overnight cultures (grown from a single colony) into fresh media, where at least 6 doublings occured and OD600 0.5-1 corresponding to 1-2*10^7 cells mL^-1^ is reached.

Cultures in the exponential growth phase (OD 0.5-1) were washed with distilled water twice and resuspended at OD 0.5 in full SD media (SD) or SD media with adenine omitted (SD ade-).

All cultures were incubated on a rotary shaker, +30°C, 180 rpm.

To demonstrate changes in optical density during starvation 96 well multimode reader Tecan Infinitie M200 with following cultivation cycle: 490 sec orbital (3,5 mm) shaking, waiting 60 s, optical density measurement at 600 nm. Alternatively, culture growth dynamics was measured with Z2 Cell and Particle Counter (Beckman Coulter).

### NMR analyses of extracellular amino acids and purines

Cell free culture media was mixed with DSS (sodium 4,4-dimethyl-4-silapentane sulfonate) in D_2_O to obtain a final DSS internal standard concentration of 1.1 mM and transferred to 5 mm NMR sample tube. NMR analysis was performed at 25°C on a 600 MHz Bruker Avance Neo spectrometer equipped with a QCI quadruple resonance cryoprobe. The noesypr1d pulse sequence was used with water suppression during the recycle delay of 10 s. The spectral width was 11.9 ppm and 128 scans were collected into 32K data points using an acquisition time of 2.3 s.

The acquired ^1^H NMR spectra were zero-filled once, and no apodization functions were applied prior to Fourier transformation. Phase and baseline corrections were applied manually. Spectra were referenced to DSS (at 0.00 ppm). The identification and quantification of sample components was performed using Chenomx NMR Suite Professional software (version 5.11, Chenomx Inc., Edmonton, AB, Canada).

### Cell morphology measurements

#### Budding index and cell size

Cell samples before and after 4 hour cultivation in media with (SD) or without adenine (SD ade-) were fixed in formaldehyde 0.5 % and examined with the optical microscope (Olympus BX51, Japan). Microphotographs (1360 x 1024 pixels) were obtained by a digital camera (Olympus DP71, Japan). Cell size and budding index were determined by microphotograph analysis in the ImageJ program.

Budding index was defined as the proportion between the number of cells with buds and total cell number. Bud was defined as a cell with cross section area less than half of mother cell size.

Cell size was determined as cell cross section area measured from the microphotographs using ImageJ program. Cells were defined as ellipses, area measured in pixels and recalculated to square micrometers (1 μm = 5,7 pixels). For each sample at least 500 cells were measured.

### Flow cytometry

Cell DNA content was determined by flow cytometry as described by Sein et al., 2018. Briefly, 0.5 mL of yeast culture was fixed in 10 mL of ice-cold 70% ethanol for at least 15 min and washed once with 50 mM citric acid. RNA was degraded using RNase A (10 μg/mL) in 50 mM citric acid overnight at 37 °C. DNA was stained with 10x SYBR Green (Invitrogen) in 50 mM citric acid for 30 min. Cells were analysed with FACSAria (Becton Dickinson, USA) device. Cell cycle distribution was analysed with Cyflogic software.

### Fermentation and metabolite flux measurements

Fermentation were done in Sartorius Q-plus fermentation system with working volume of 0.3 L. Gas flow 0.25 L * min^-1^, mixing rate 400 rpm, media pH was set to pH 5.5.

Biomass concentration was determined as absorbance in 590 nm (WPA Colorimeter Colourwave CO7500, Biochrom, UK). Following coefficient to convert absorbance units to dry weight was used 1OD_590_ = 0,278 g*L^-1^. Carbon dioxide evolution was recorded by an exhaust gas analyser (InforsHT) in parallel to harvesting of metabolite samples.

Extracellular glucose, ethanol, acetate and glycerol contents were measured simultaneously by Agilent 1100 HPLC system with a Shodex Asahipak SH1011 column. Glucose, acetate, glycerol and ethanol were quantified with a refractive index detector (RI detector RID G1362A). The flow rate of the mobile phase (0.01 N H_2_SO_4_) was 0.6 ml min^-1^; the sample injection volume was 5 μL.

Biomass from fermentations were centrifuged and Intracellular nucleotide pools extracted via cold methanol extraction. ATP, ADP and AMP were quantified by HPLC-MS-TOF analyses, as described in (Valgepea et al, 2012).

### FTIR data

For cell macromolecular content analysis FTIR spectroscopy was used as described in Grube et al. 2002. 2 mL of cells (OD1-4) were harvested by centrifugation and washed 3 times with distilled water. Cell pellet was diluted by 50 uL of distilled water, samples were spotted on 96 field spot-plates. Absorbance data were recorded by Vertex 70 with microplate extender HTS XT, interval 4000 - 600 cm^-1^, resolution 4 cm^-1^. For data collection and control OPUS/LAB, OPUS 6.5 software was used.

### Cell *carbohydrate* extraction and quantification

Fractional cell polysaccharide purification for quantitative assays was done as described in Stewart, 1975. Total saccharide content of each fraction was determined by anthrone assay, results were expressed as glucose equivalents mg*gDW^-1^ biomass. (Dubois et al., 1956).

### Transcriptomics

Total yeast RNA after 4 hour cultivation in synthetic dextrose (SD) or SD media with adenine omitted (SDade-) were isolated with RiboPure™ RNA Purification Kit (Thermo scientific). Yeast transcriptome was analysed using MiSeq (Illumina) NGS data analysis. See supplementary material 4 for details.

Expression data set was submitted in European Nucleotide Archive (ENA) database, accession no. PRJEB40525.

### Sublethal stresses

Cells were grown in SD media until exponential phase, washed with distilled water twice and inoculated in SD or SD ade-media with cell density 1*10^7^ cells*mL^-1^. After 4h of incubation cells were harvested by centrifugation, washed with distilled water once and aliquoted in 1 mL, OD600=1 (corresponds to 2*10^7^ cells*mL^-1^). 3 aliquots were exposed to each stress. Thermal stress - cells were kept in 53°C for 10 min. Oxidative stress - cells were incubated in 10 mM H_2_O_2_ for 50 minutes, afterwards washed with distilled water. Desiccation - cells were sedimented by centrifugation, supernatant removed and pellet air dried in the exicator, 30°C for 6 hours. After drying, distilled water was added to resuspend cells.

After all stress treatments cells were serially diluted and dilutions spotted on YPD plates to assess CFU*mL^-1^. To check for cell losses during washing steps OD of suspension was measured and CFU*mL^-1^ corrected for OD value. Survival is expressed as % assuming that OD600=1 corresponds to 2*10^7^ cells *mL^-1^.

To test weak acid stress resistance cells were spotted on YPD plates supplemented with 0.1 M acetic acid, with pH of agar media set to pH 4.5 (Martynova et al., 2016).

## Results

To characterise global changes initiated by purine starvation, we constructed *ade8* knockout in laboratory yeast strain CEN.PK2-1D background (*ade8* strain). To dissect physiological effects imposed by purine starvation in the CEN.PK2-1D background, we cultivated *ade8* strain in SD media with all necessary auxotrophic supplements present in surplus (here and rest of the text called SD media) and in the media with adenine omitted (SDade- media).

### Growth of CEN.PK2-1D ade8

CEN.PK2-1D strain is often used in laboratory experiments and contains several auxotrophic markers. These are histidine (*his3*), leucine (*leu2*), tryptophan (*trp1*), uracil (*ura3-52).* We introduced additional purine auxotrophy by “clean” (antibiotic marker free) *ade8* knockout as suggested by Sadowski et al. 2008. Lack of any single supplement necessary to complement the metabolic needs of this strain leads to cease of growth (Fig 1A). When exponentially growing *CEN.PK2-1D ade8* cells were washed of media and inoculated in SDade- media, cells show increase in cell numbers in first two hours, but afterwards cell number mL^-1^ stays stable (Fig 1B). For most of our experiments we chose cells that have been starved in SDade- for 4h, as that is the time when ade-specific phenotype appears. To be sure, that we are characterising *ade8* mutation as only factor influencing cell phenotype, we made sure that all other auxotrophic agents are still in media after 4h of purine starvation (Fig 1C)

**Figure 1.**
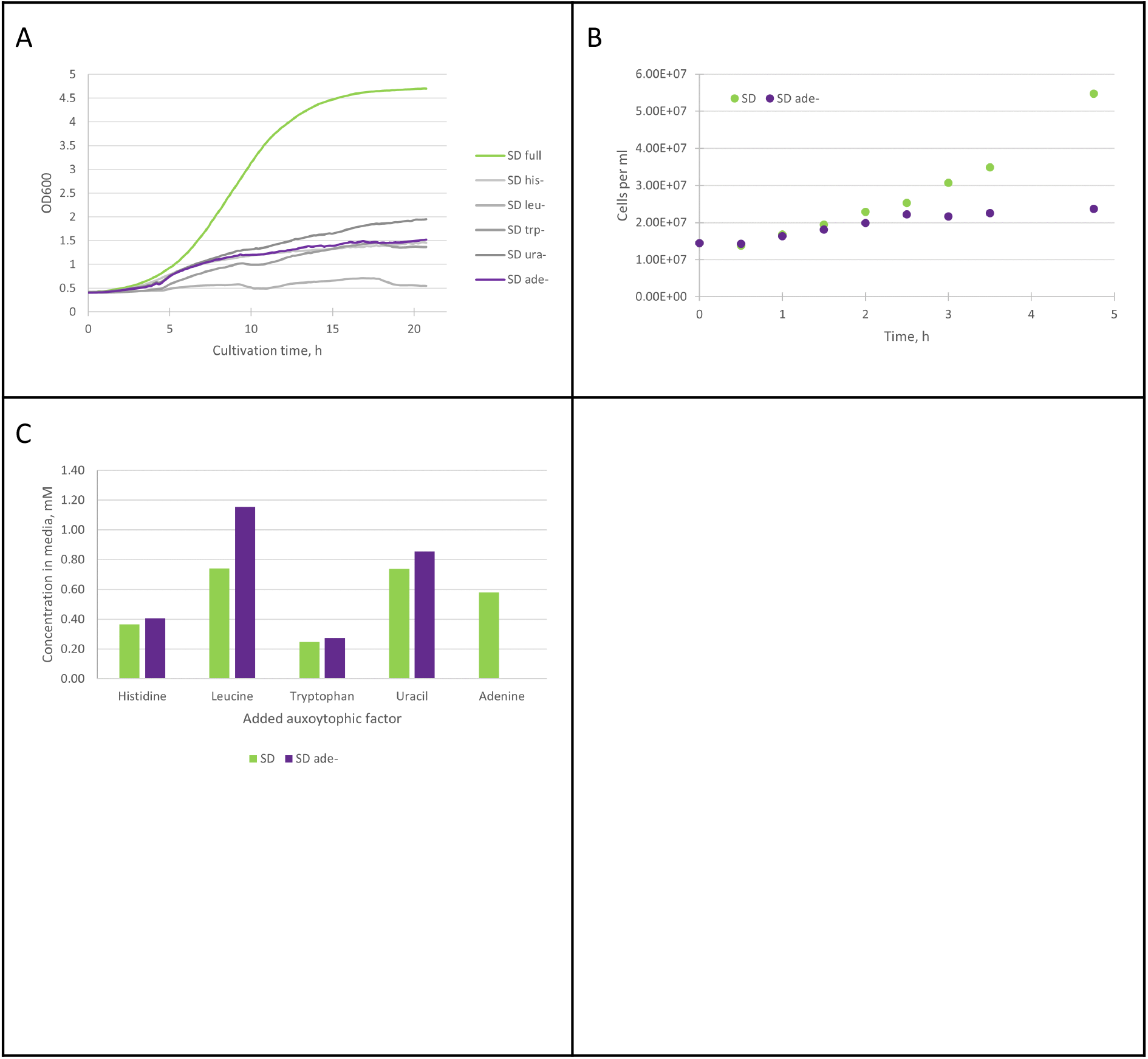
**A** Growth of *ade8* (CEN.PK2-1D MAT alpha *his3D1; leu2-3_112; ura3-52; trp1-289; MAL2-8c; SUC2 ade8Δ0*) strain in SD media with all auxotrophic factors added in surplus or one of auxotrophic factors omitted to starve cells for that particular nutrient. A slight increase in optical density can be observed with most starvations **1B** Cell number per mL during first 4 hours of growth in SD media or starvation for adenine. It can be seen that cells stop increasing in numbers after 2h of purine starvation **1C** Amount of auxotrophic factors in growth media after 4 hours of *ade8* cultivation. In purine starvation media all other auxotrophic factors except adenine are still in surplus

Therefore, we conclude that our media composition can induce starvation for adenine (purine) specifically and physiological effects observed thereof are solely due to lack of external purine supply.

## Cell morphology and cell cycle

### Purine starvation stops cell cycle

Although cell number*mL^-1^ did not increase after the 2nd hour of *ade8* cultivation in purine starvation media, elevation of optical density over time was observed (compare Figure 1A and B). This led us to hypothesize that specific changes in cell morphology occurs, thus increasing light dissipation and account for OD increase during *ade8* purine starvation. We quantified the budding index, analysed DNA content of the cell using FACS and measured cross sections of cells of the *ade8* grown in SD and SDade- media.

Budding index is an indicator of culture progression through the cell cycle. We defined the budding index as the ratio [%] of the number of cells with buds and the total number of cells, see figure 2A.

**Figure 2.**
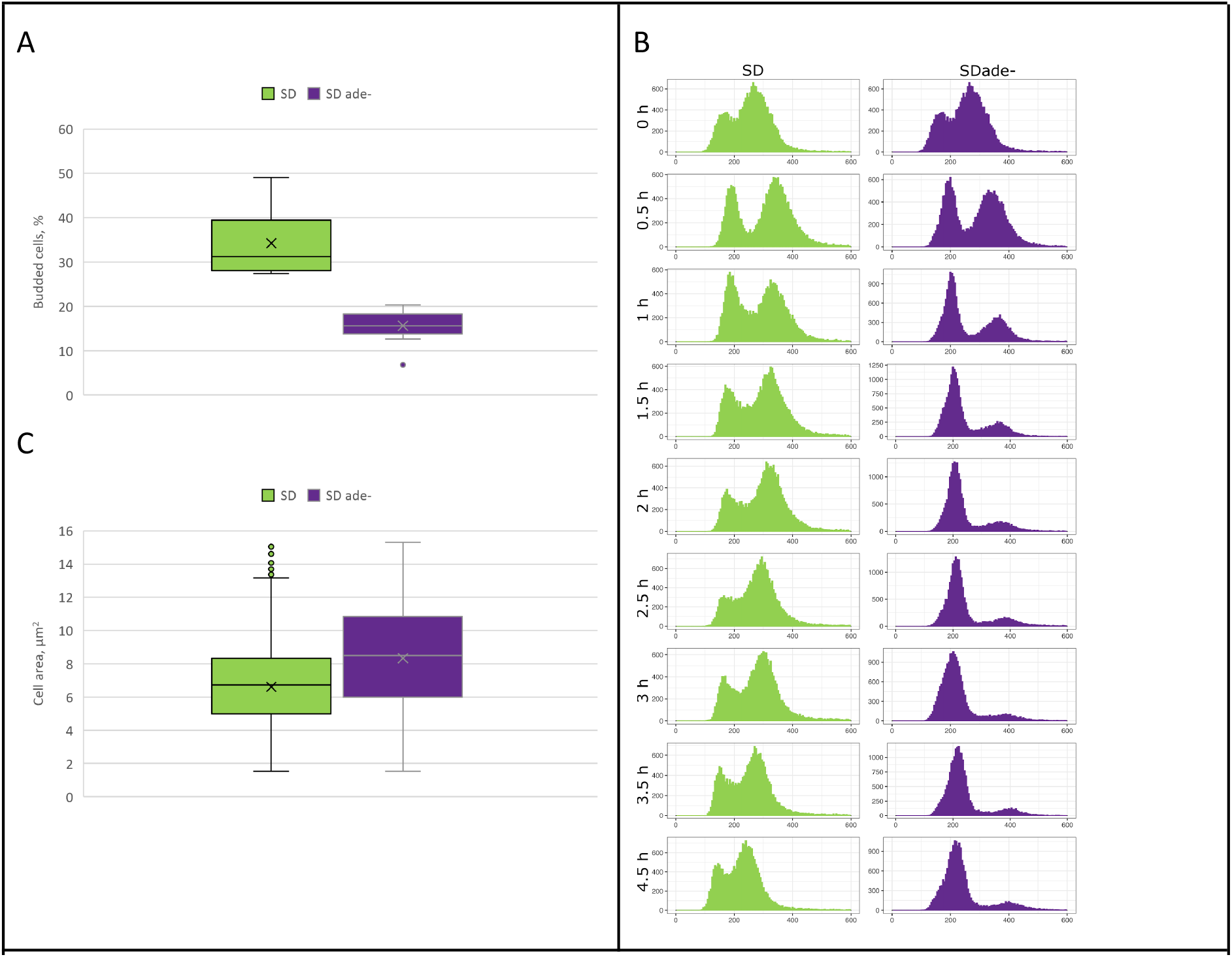
*ade8* cell morphology changes when grown in full or adenine deficient media **2A** Cell budding index. Data from 200 cells analysed from microscopic images **2B** *ade8* strain cell DNA copy dynamics over time. in SD and SDade- media. **2C**. Cell size analyses as determined by area of cell cross-section in microscopic images. Data from at least 500 cells from each cultivation.

We observed that budding index is approximately 30% when cells are cultivated in SD media, while in case of purine starvation media it significantly decreases (down to 15%).

To complement budding index data and test DNA content in the cell of a population grown in SD or SDade- media, we performed FACS analyses. We found that in the case of purine starvation culture becomes enriched with cells harbouring N copies of DNA per cell. Moreover, this occurs within the first two hours of cultivation. After 4 hours of cultivation *ade8* cell population when cultivated in SD media contains both N and 2N DNA copies per cell while cells cultivated in SDade- contain mostly N copies of DNA (compare DNA peak dynamics in SD and SDade- media, Fig 2B).

We tested if cell size changes could contribute to OD increase during purine starvation. We measured the average cell cross section after 4 h cultivation in SD and SDade- media. Indeed, purine starved *ade8* cells are larger than cells growing in SD media, see in Figure 2C. Therefore we conclude that purine starvation leads to a drop in budding index which is accompanied by increase of cell population with cells with N copies of DNA and significant increase in cell size (as demonstrated by increase in cell cross section). These morphological markers demonstrate that purine starved cells are morphologically different from cells growing in SD media.

## Biochemistry of purine starvation

Changes in culture growth parameters (optical density or cell concentration) are probably the most obvious phenotypic markers for auxotrophic starvation. To further dissect purine starvation effects, we measured several metabolic markers when *ade8* was cultivated in SD or SDde- media. We measured glucose consumption and production of major carbon metabolites (ethanol, CO_2_, glycerol, acetate and biomass) as well as we determined intracellular ATP, ADP and AMP concentrations.

Specific growth rate [h^-1^] of *ade8* in SD media was 0.4 and in SDade- media 0.15. Glucose specific uptake q [mCmol * g DW^-1^ * h^-1^] during purine starvation was two times slower than in cells in SD media: 43 +/- 3.6 and 81+/-5.4 mCMol *gDW^-1^*h^-1^ respectively. Meanwhile specific CO_2_ production rate was 5-6 times higher in SD media than in SDade- media (see figure 3B). Since glucose was the sole carbon source and we could account for more than 90% of carbon in total, we calculated flux distribution as carbon % of glucose consumed for *ade8* cultivated in SD or SDade- (See figure 3B and raw flux data in supplementary materials 1).

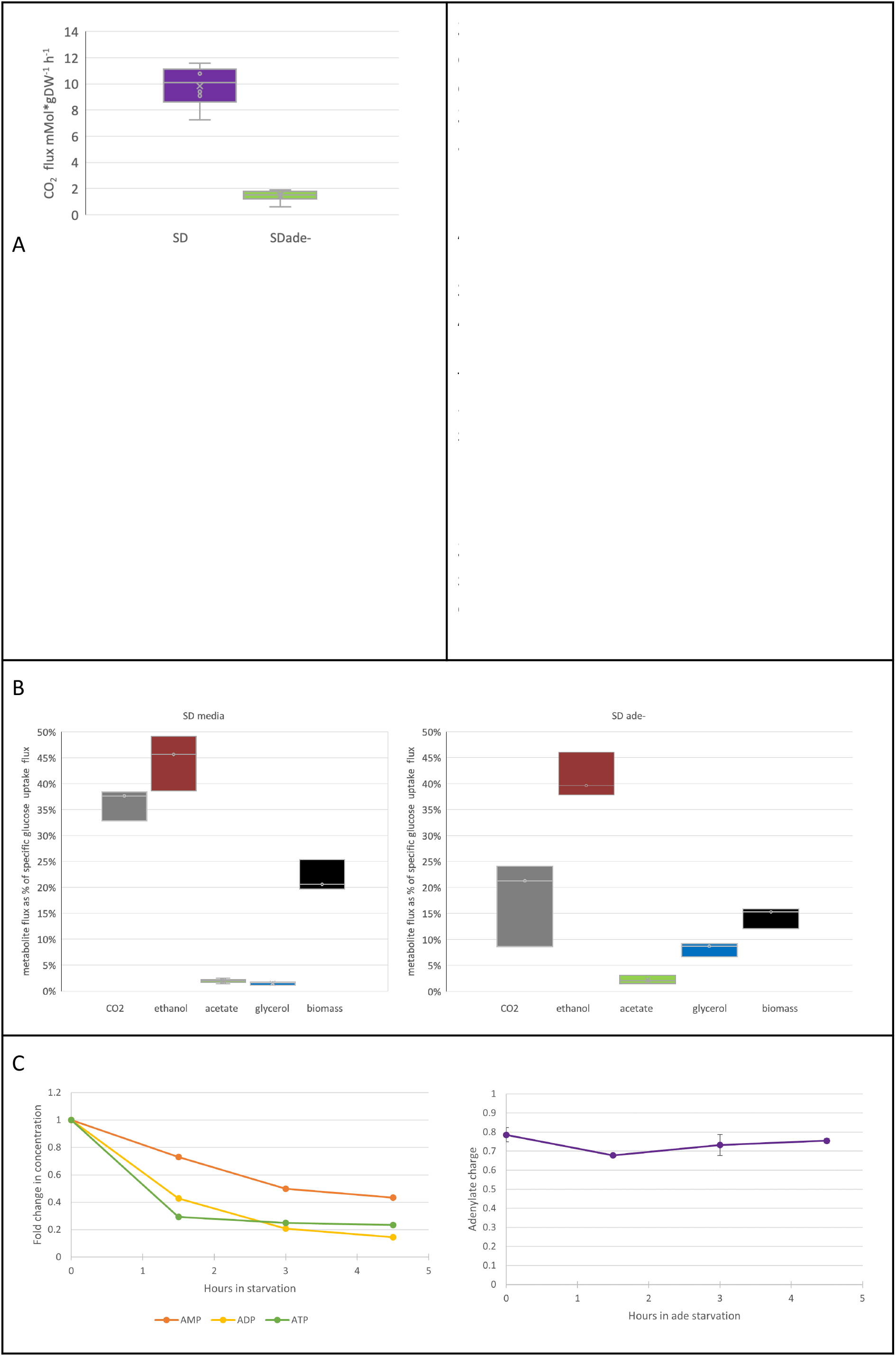
**3A** Specific CO_2_ production carbon flux distribution in the case of *ade8* strain cultivation in SD or SDade- media **3B** Carbon flux distribution in the case of ade8 strain cultivation in SD full media or in media with adenine omitted Average CO_2_ flux was measured as produced [mM *gDW-1*h-1], from SD and SDade- media. All cultivations were performed in batch mode, in 400 mL Sartorius Qplus fermentation system. Starting volume was 300 mL. CO_2_ was measured by infrared CO_2_ sensor (InforsHT) Box plot depicts standard deviations, error bars - min and max values from three independent bioreactors. **3C Left,** Changes of AXP amount in ade starved *ade8* cells. **3C Right** Adenylate charge during purine starvation.

When cell growth is suspended due to lack of essential metabolite (purine), not only main carbon fluxes are affected (fig 3A,B), but also concentrations of intracellular purine nucleotides. We measured concentration of purine containing moieties (ATP, ADP and AMP) to find out if intracellular concentration of these molecules changes if external supply for their precursor adenine is diminished (see Figure 3C).

Indeed, already 1.5 h after SD media shift to SDade-, intracellular concentration of ADP and ATP dropped more than half of initial intracellular concentration (Figure 3C). Interestingly, while intracellular concentration of ATP, ADP and AMP dropped significantly during purine starvation, energy charge throughout purine starvation remained almost constant. There is a slight drop in the beginning of starvation, but adenylate charge reaches pre-starvation levels in the cells after that. It should be kept in mind that in the first two hours of purine starvation cells are still proliferating (see Fig 1B).

Although *ade8* growth in SDade- media is stopped, cells continue to metabolise glucose. However, specific glucose uptake dropped significantly from 81+/- 5 to 43 +/- 3 (mCmol *g DW*h^-1^). We think that it is related to decrease of intracellular adenine nucleotide concentrations (Fig 3C) which does not allow rapid glucose metabolism (Larsson et al., 2000). Also distribution of other carbon fluxes was altered. In SD media most of energetic needs seems to be fulfilled with the help of fermentation, still, there is also CO_2_ production that is not coming from ethanol production, pointing to the involvement of mitochondrial activity and respiro-fermentative growth. In SDade- conditions all CO_2_ can be attributed to ethanol production. Interestingly, glycerol production is significantly increased.

It seems that instead of biomass a significant amount of carbon has been redirected to glycerol synthesis, which points to an increase of cell glycerol and/ or lipid content as glycerol is the backbone of triacylglycerols (TAG).

We checked if macromolecule content of the biomass is affected when cells are cultivated in SD media or purine starved. Relative amounts of proteins and nucleic acids decrease, while carbohydrates and lipids increase (See Fig 4A).

**Figure 4.**
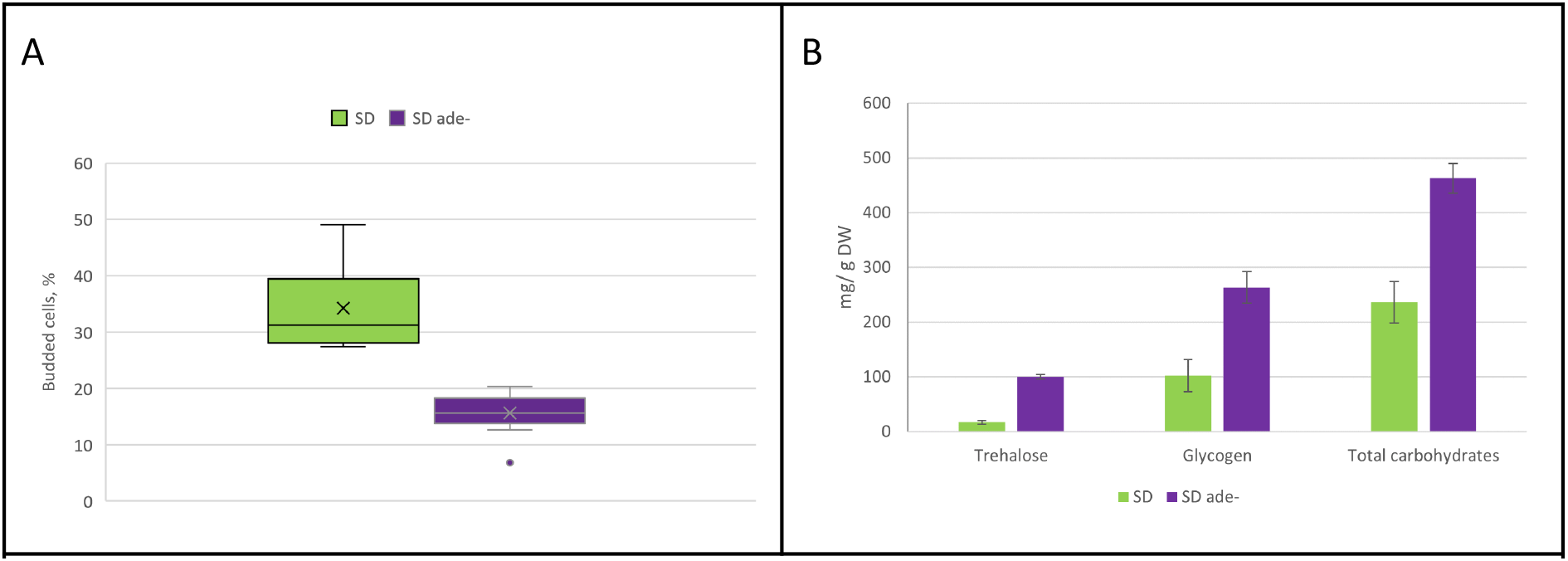
**A** Distribution of macromolecules in cell biomass as assessed by FTIR. Data is average from two biological replicates, error bars show standard deviation among technical replicates. **4B** Amount of carbohydrates in biomass assessed by the anthrone method. Data shown is average from three biological replicates. Error bars - standard deviation of biological replicates.

We extracted fractions of main budding yeast reserve carbohydrates - trehalose and glycogen (Stewart, 1975) and quantified saccharide content of each fraction by anthrone method (Dubois, 1956). Results show that during 4h of purine starvation *ade8* cells accumulate 100 mg trehalose and 263 mg glycogen per g DW. While cells that are growing in SD media had 17 mg and 102 mg of trehalose and glycogen per gDW respectively. We also measured concentration of other cellular carbohydrates that make most of yeast biomass - mannans and beta-glucans. It can be seen that an increase in carbohydrate fraction during purine starvation is due to accumulation of reserve carbohydrates, while amount of structural carbohydrates does not change (Fig 4B). Purine starved yeast biomass have accumulated more reserve carbohydrates by 244 mg*gDW^-1^ and total carbohydrate content increased by 227 mg*gDW^-1^, which correlates to an increase of carbohydrate fraction in biomass macromolecular composition. Carbohydrate fraction in purine starved cell biomass almost doubled, as revealed by FTIR analysis, see Fig 4A.

To verify biomass macromolecular composition data (Fig 4A), we estimated RNA content of the cells. We assessed RNA quality (Qiaxcell capillary electrophoresis) before expression analysis by comparing 18S and 28S rRNA ratio. The ratio between 18S / 28S did not change (whether it was SD or ade-sample), therefore RNA quality was good, no degradation was observed. At the same time, the amount of extracted RNA per unit of biomass was 3 times smaller in case of adenine starved cells, thus reinforcing biomass macromolecular content obtained by FTIR - decrease of nucleic acid amount during purine starvation.

## Sublethal stress resistance

Carbon flux distribution away from energy production towards storage metabolites, increased glycerol production, decrease in glucose uptake and decrease of intracellular adenine nucleotide content, indicate that energy and carbon in the case of purine starvation might be redirected to other functions, away from biomass synthesis and growth.

Stress resistance is an important phenotypic feature of microbial cells. It is known that general environment stress resistance (ESR) phenotype is induced in slowly growing, stationary or quiescent cells (reviewed in Gasch & Werner-Washburne, 2020). Previously it has been shown that methionine auxotrophic starvation can lead to elevated stress resistance (Petti, et al., 2011). To test if cultivation in purine starvation media affects culture stress resistance we tested cell viability after exposure to harsh environmental stresses (sublethal stress resistance). We tested if purine starved cells have higher resistance to short thermic or oxidative stress as well as weak acid stress (growth on plates containing acetic acid at pH 4.5, to see long term stress resistance). Additionally we tested desiccation tolerance as an example of multicomponent stressor. The time of exposure of each stress was adjusted so, that viability of exponentially growing cells to be 10%.

We observed that survival of purine starved cells was more than 10 fold higher than for cell population growing in SD media in all sublethal stresses tested, see Fig 5A.

**Figure 5.**
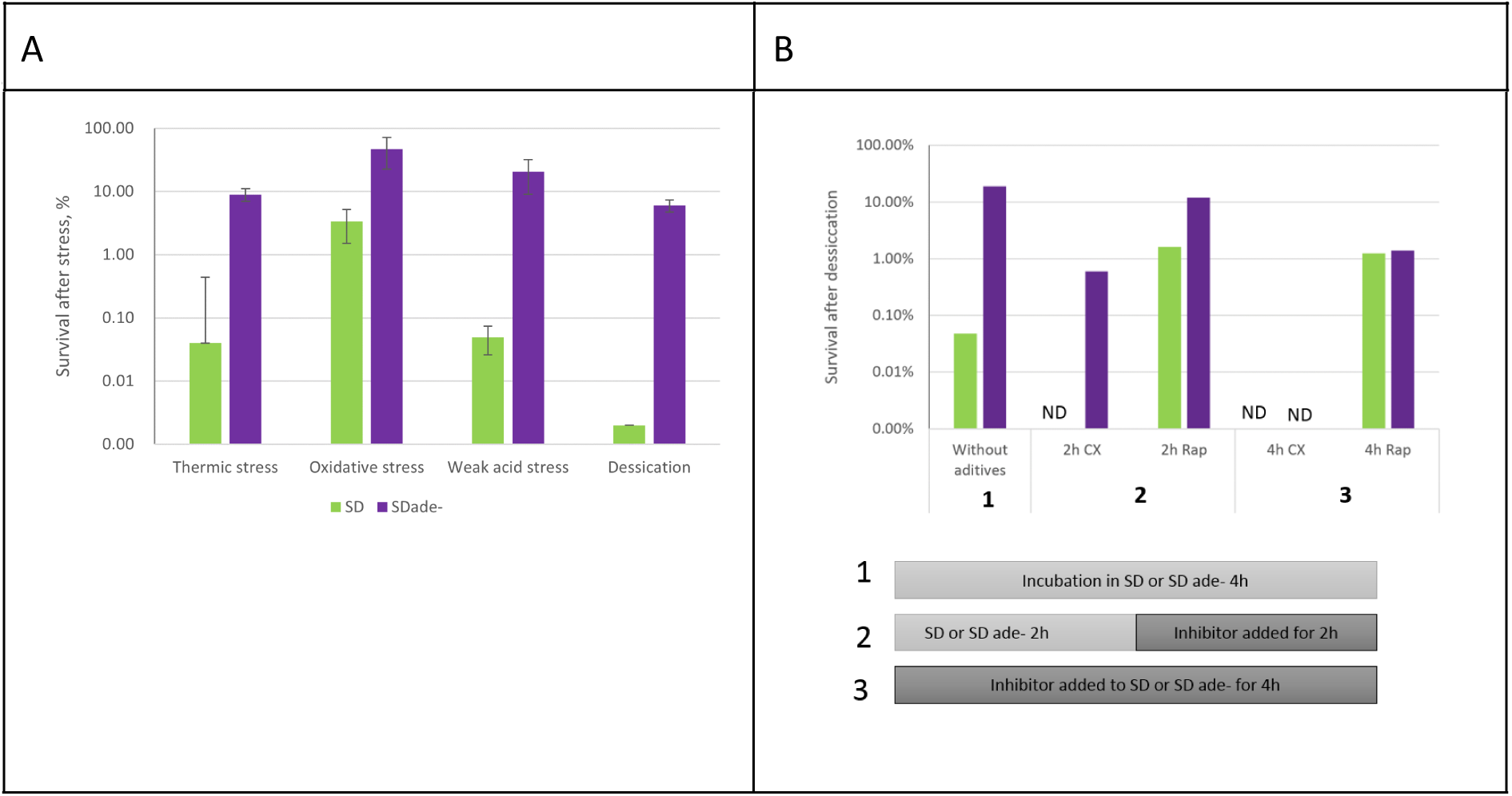
*ade8* strain stress resistance after cultivation in SD or SDade- **5A** Cell survival after exposure to sublethal stress conditions. *ade8* cells cultivated in SD and SDade- were exposed to thermic stress (53°C, 10 min), oxidative stress (10 mM H_2_O_2_, 50 min) weak acid stress (plated on YPD with 0,1 M acetic acid, pH 4.5) or dessicated (30°C for 6h) after stress cfu*mL^-1^ OD^-1^ was assessed by plating on YPD plates. **5B** Top panel cell survival after desiccation and addition of cyclohexamide or rapamycin during starvation. Bottom panel - setup of experiment. Cells were treated with cyclohexamide or rapamycin in various time points during purine starvation and then desiccated (30°C for 6h) and plating was done as in other stress assays. **NB!** The scale of y axis in both graphs is exponential.

We wanted to understand if desiccation tolerance is a phenotype evolving immediately after shift to starvation media or some “adaptive reactions’’ occur while cells are starving for purine. To test this development of stress resistance phenotype, we used desiccation assay, since this treatment gave the most distinct signal for cells cultivated in SDade- and SD media. To determine that, we added cyclohexaminde - translation inhibitor to cells either in the beginning of starvation or 2 hours after start of starvation, when active cell proliferation has ceased. The results are depicted in Fig 5B. It can be seen that translation inhibition during auxotrophic starvation lowers desiccation tolerance after starvation. Therefore, we conclude, that indeed, stress resistant phenotype develops during purine starvation via new protein production.

Our results demonstrate that purine starvation preconditions cells to become stress resistant. However, it is not known how cell coordinates lack of purines to these massive phenotypical changes. Other authors have shown that desiccation tolerance can be TOR pathway dependent (Welch et al., 2013). This fact together with the decrease of RNA content, led us to inquire if TOR signaling system mediates purine starvation to development of specific phenotype. We used desiccation tolerance as a marker of purine starvation phenotype and compared desiccation tolerance of purine starved cells and SD media grown cells, adding rapamycin during incubation in an experiment setup similar to cyclohexamide assay. Addition of rapamycin did not affect desiccation tolerance significantly (see Figure 5B). For purine starved cells, if no rapamycin was added, 19% cells were desiccation tolerant, after 2 h rapamycin treatment 12%, and after 4h of rapamycin treatment 1% of cells survived desiccation. Addition of rapamycin to SD grown cells increased their desiccation tolerance, but not to a degree of purine starved ones: no rapamycin 0.05%, 2h rapamycin 1%, 4h rapamycin 1% (see Figure 5B). Since purine starved cells exhibit desiccation tolerance 10-20x times higher than rapamycin treated, we conclude that the TOR system might be involved in purine starvation signaling, but there are additional systems in play.

## Transcriptomic response

Results from cyclohexamide assay (Figure 5B) shows that stress resistance is driven by gene expression, not just caused by disturbances in intracellular metabolites. To assess gene expression changes over purine starvation, we performed RNAseq.

RNA expression data was gathered from cells that have spent 4h either in SD or adenine deficient media. When comparing expression data of *ade8* cells cultivated in SDade- or SD, 455 upregulated genes (more expressed in SDade- conditions) and 244 downregulated genes (more expressed in SD) were identified. Also 6362 non-significant gene expression changes were noted (−2<logFC<2). Among the top 20 most upregulated transcripts we found genes coding for stress resistance proteins (*SIP18* and it’s paralogs, *DDR2, HSP12, HSP26*), stationary phase response proteins (*SPG4* and *SPG1*) and carbohydrate metabolism (*HXT5, HXT6, TKL2, GND2*). Interestingly, expression of several spore related genes were also upregulated, for example *SPS100.* For full gene list see Supplementary material 3.

Most prominent downregulation was observed in expression of various various tRNA and protein genes related to the transcription process. This is also demonstrated by performing GO term enrichment search. Results are depicted in Fig 6. Metabolic process enrichment terms group in two clusters that correspond to up and downregulated genes respectively (Figure 6A). Genes that are the most represented in our data set are connected with downregulated genes and are connected with the translation process. From upregulated genes ox-red processes and carbohydrate metabolism are GO terms that are most enriched in ade-starved cells. Table with all gene GO terms and their expression statistics can be found in supplementary materials 2.

**Figure 6.**
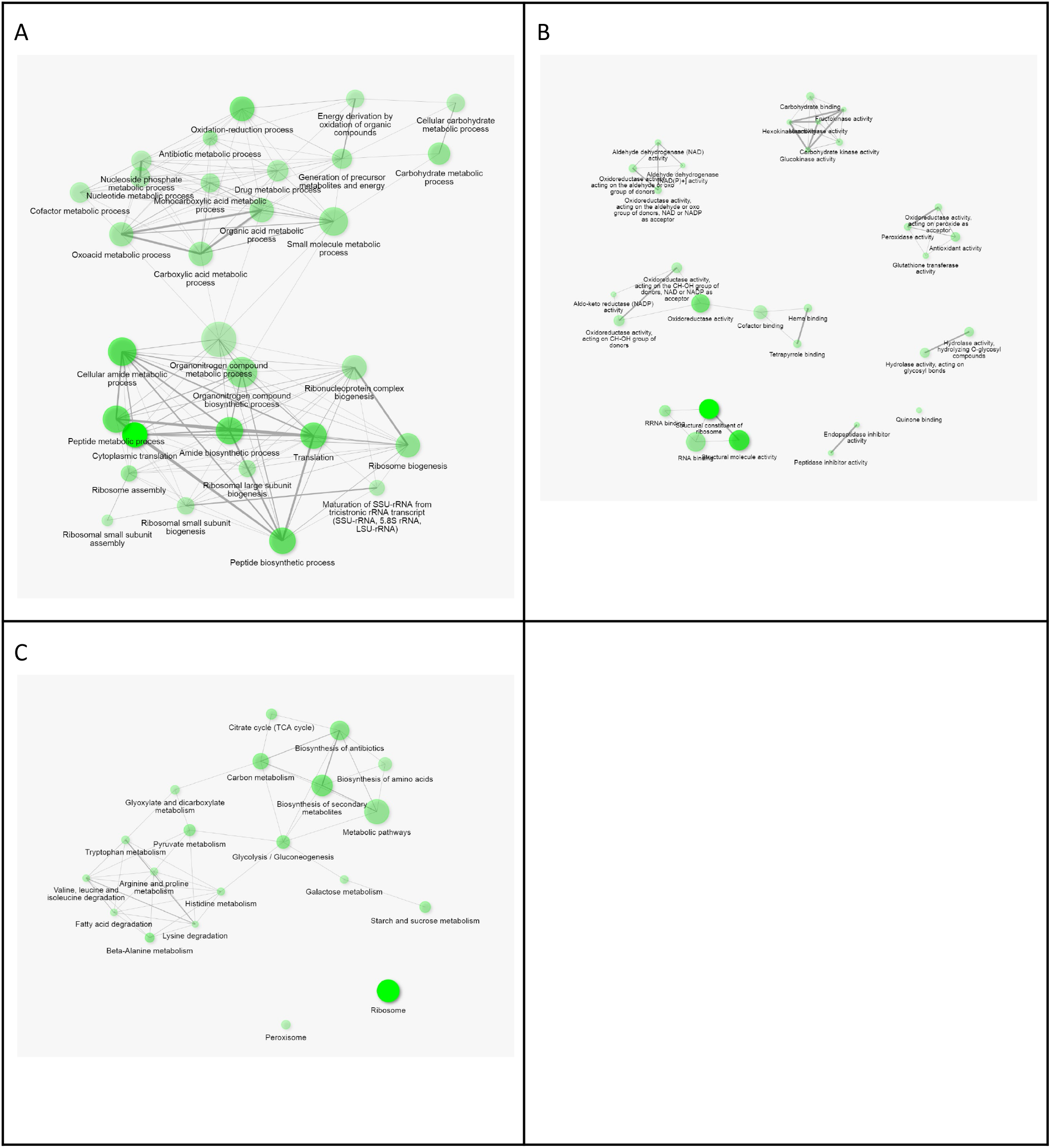
Gene pathway enrichment analysis. From transcription results genes that were at least two fold up or downregulated in SDade- cells when compared to SD cells cells were selected. For those genes enrichment analysis was made and plotted using ShinyGO v0.61. Plot also shows the relationship between enriched pathways. Two pathways (nodes) are connected if they share 20% (default) or more genes. Darker nodes are more significantly enriched gene sets. Bigger nodes represent larger gene sets. Thicker edges represent more overlapped genes. **6A** Enrichment in GO metabolic process terms **6B** Enrichment in GO molecular functions **6C** Enrichment in KEGG pathway terms

Also if all upregulated and downregulated genes are plotted on TheCellMap.org (see Fig 7), where interactions among genes are visualised, it can be seen that down regulated genes group strongly around ribosome biogenesis cluster, but upregulated genes do not show such a specific grouping.

**Figure 7.**
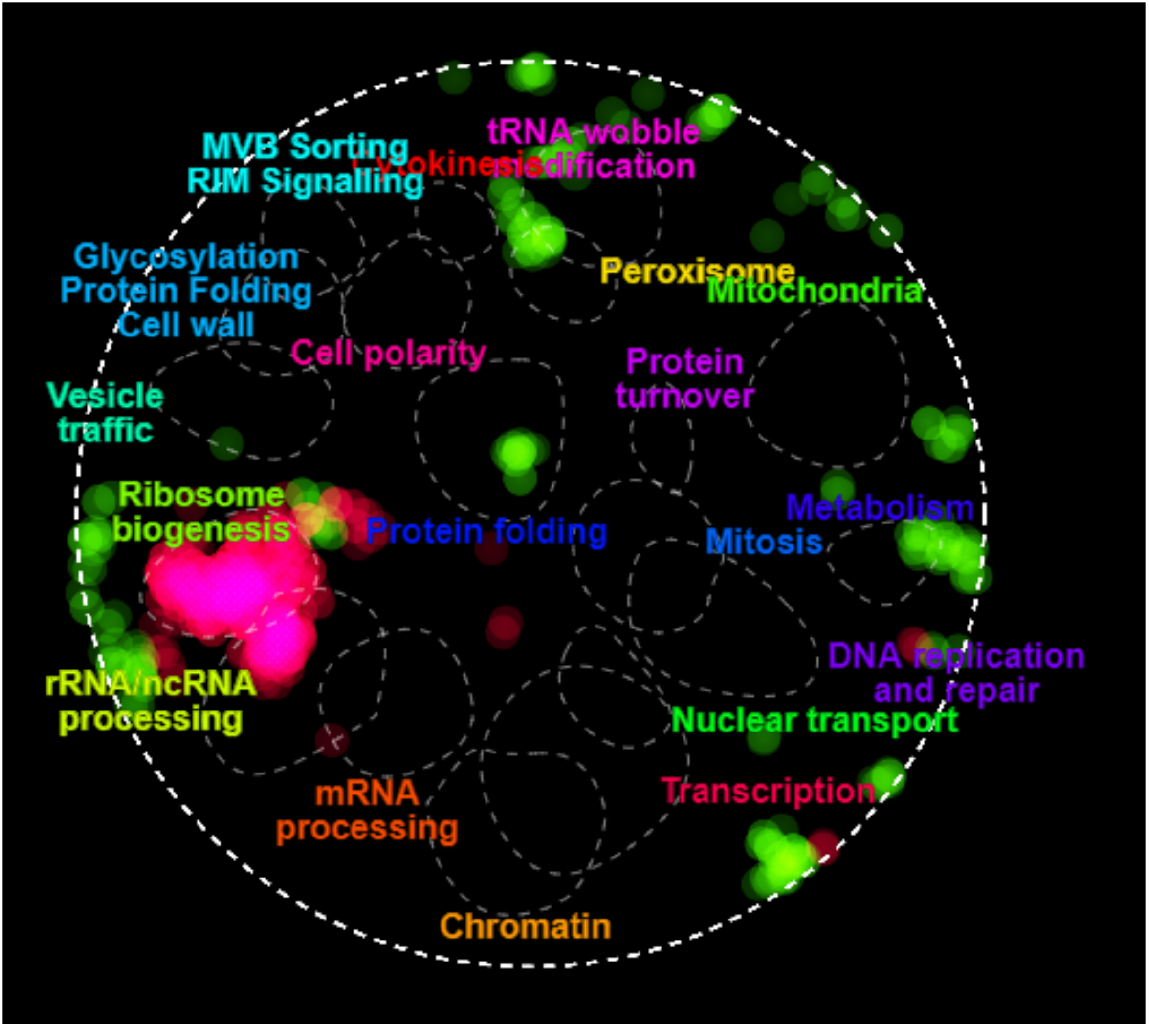
Genes that are upregulated - green and downregulated - pink in purine starved cells when compared to cells cultivated in SD media. Plotted with cellmap.org.

## Discussion

Nowadays it is easy to knock out practically every gene in many model organisms. However, the phenotypic effects caused by such a deletion cannot always be extrapolated from the knowledge of encoded protein function alone. This is partially due to the fact that metabolism is highly interconnected and metabolic intermediates once synthesized in one pathway are consumed in other pathways. Therefore deletion of a gene coding for enzyme in a pathway producing metabolite of high demand might potentially affect many reactions in seemingly unrelated pathways and subsequently – number of different organism functions. This has been shown in the case of *zwf1* knockout strain, where oxidative tolerance is severely affected due to defect in NADP/H recycling (Campbell *et al.*, 2016). Our results imply that a gene knockout (*ade8*) of purine synthesis pathway is another example where deletion has indeed global phenotypic effects. Moreover, to ensure survival, strong interaction between organism and its environment occurs. Therefore different environmental conditions and gene/s interact and might lead to different phenotypes. In *ade8* strain most striking phenotypic effects develop during purine starvation. This is a seemingly specific environmental niche, yet purine auxotrophic starvation might set rapidly and affect cell phenotype (even within 2 hours) when media purine is depleted.

### Intracellular signalisation of purine starvation

We observed that cells have a lower amount of rRNA after 4h of starvation. After the beginning of purine starvation most yeast cells finish the DNA synthesis phase and halt budding; putative arrest in G1 can be observed (see Fig 1B and 2B). Most of cell’s RNA is ribosomal RNA (rRNA), therefore by degrading rRNA it would be possible to free nucleotides required for DNA synthesis. At the same time analysing expression data, expression of ribonucleotide reductase *RNR2* was not significantly altered (*RNR2* in SD media logFC=2.04 and after 4h in ade- media log FC=1.64).

Pelletier and colleagues used carcinoma cell lines to see how nucleotide depletion affects the cell cycle. Their research shows that when purine levels drop, ribosome assembly is delayed, as nucleotides are needed for RNA. This causes failure of cell cycle checkpoint and p21 accumulation, arresting cells in G1 phase (Pelletier et al., 2020). Overexpression of *S. cerevisiae* p21 analog *CIP1* also causes arrest in G1 phase. Hyperosmotic stress also causes yeast cells to activate *CIP1* via Msn2/4p thus delaying cell cycle (Chang et al, 2017). When analysing our expression data with YEASTRACT tool (Abdulrehman et al., 2011), Msn2p and Msn4p transcription factors were suggested as one of possible regulators (see full list with transcription factors in Supp2), also our previous research shows that Msn2p and Msn4p are involved in purine starvation elicited stress resistance (Ozolina et al, 2017). Thus involvement of Cip1p in purine starvation sensing should be considered and researched in more detail.

Hoxhaj and colleagues explored how HeLa cells react to purine analog. They notice similar patterns as we do - intracellular concentration of AMP, ADP and ATP decreases, but cellular adenylate charge is kept constant. They propose that purines are sensed with the help of mTOR signaling system, where TSC complex would be responsible for sensing lack of adenine and inhibiting mTOR pathway further on (Hoxhaj, et al. 2017).

Our expression analysis shows downregulation of translation machinery that is consistent with involvement of TOR signalling, at the same time it is necessary to note that *Saccharomyces cerevisiae* lacks TSC complex (Dibble, Manning, 2010). Additionally, we compared the effect of TOR inhibition by rapamycin with purine starvation on yeast desiccation tolerance (see Fig 5B). We observed that purine starvation induces increased stress desiccation resistance much higher than rapamycin alone. Moreover, additional rapamycin together with purine starvation did not increase desiccation tolerance. Therefore, if TOR system is involved in purine sensing in *S. cerevisiae*, it receives a signal from an alternative signal transduction pathway, upstream of canonical TOR protein Frp1p (Koltin et al., 1991).

### Purine starvation affects adenylate pool, not charge

Exponentially growing cells devote significant energy resources for biosynthesis of new cells. When cell growth is halted, a significant amount of energy is rerouted for “maintenance reactions” (Vos et al. 2016). Most purines are used in the synthesis of nucleic acids, hence the need for these moieties in nondividing cells is reduced. Also most purine biosynthesis genes are downregulated at the transcriptional level upon entry into stationary phase (reviewed in Ljungdahl & Daignan-Fornier, 2012).

Purine biosynthesis is regulated by ATP levels via Ade4p activity and downstream metabolites AICAR and SAICAR activating Bas1p and Bas2p transcription factors. Bas1/2p are responsible not only for activation of purine biosynthesis genes, but also all pathways connected with purine biosynthesis - histidine, glutamine and one carbon metabolism. Bas2p (also known as Pho2p) is a regulator of phosphate metabolism (Ljungdahl & Daignan-Fornier, 2012). Drop in intracellular ATP concentration in WT would activate Ade4p; additionally, in the presence of AICAR and SAICAR Bas1/2p activate other genes of *de novo* purine synthesis pathway. However, *ade8* strain strain used in this study possesses *ade8* and *his3* mutation, therefore AICAR and SAICAR cannot be synthesised via purine nor histidine biosynthesis pathways. Thus in our *ade8* strain during purine starvation no “traditional” signaling via Ade4 and Bas1/2p is not possible; therefore, response to purine shortage was sensed and signalled through other means.

ATP, ADP and AMP form the main portion of cell adenylates. Among them ATP is the most abundant free nucleotide in the cell. We saw that during purine starvation although the total intracellular amount of the AXP pool decreases, adenylate charge is maintained almost unchanged. Therefore, we think that active processes to homeostase adenylate charge are present. This is in accordance with findings from other experiments where knockout of purine metabolism genes does not affect adenylate charge. Specifically, adenylate charge in knockouts of adenylate kinase *adk1* nor AMP deaminase *amd1* is not affected (Gauthier *et al.* 2008; Saint-Marc *et al.* 2009). Adenylate level regulates many metabolic enzymes and it could be that a drop in AXP level itself acts as a specific intracellular signal for yeast cells. For example, purine auxotrophic parasite *Leishmania spp.* senses perturbation in nucleotide pools and it seems, that decrease in purine nucleotide pools signals the need to switch on the stress resistant phenotype. Indeed, starvation for purines ensures survival of *Leishmania donovani* for up to 50 days in purine scarce environment (Martin et al., 2016).

### Purine starvation affect transcription factors

To understand if purine starvation initiates a specific transcriptional pattern, we looked for which transcription factors most probably are responsible for the gene transcription pattern we observe. Also we compared our data from SDade- media with publicly available datasets of nitrogen starvation and stationary phase yeasts.

First, we saw that *ade8* cells during purine starvation do not stop their metabolism, instead, they actively produce new proteins and attain stress resistance phenotype. Desiccation tolerance is abolished if cells are purine starved and cyclohexamide treated for 4 hours. Some viability (0.6 %) is retained after 2h of cyclohexamide in ade-media (Fig 5B). Thus we conclude that during the first two hours of starvation some cell signalling processes are started that influence survival during desiccation. For these processes to take effect protein translation is required. Rapamycine addition to cells at the beginning of purine starvation results in 1% survival, while if cells were in SDade- media for 2h before addition of rapamycin, desiccation tolerance is increased by an order of magnitude. Therefore, it seems that global transcriptional response (upstream/ parallel to TOR) would drive development of stress resistance phenotype during purine starvation rather than TOR pathway alone.

When performing search for transcription factors which might be associated with gene transcription patterns during purine starvation with the YEASTRACT tool (Abdulrehman et al., 2011), we observed strong link with Dal82p and Gln3p typically associated with regulating nitrogen source dependent gene transcription. At the same time purine starvation transcriptional response was associated with other transcription factors connected with diauxic shift and carbon source management: Yap7p, Gis1p, Mig1p, Hsf1p, Hap4p, Adr1p.

We compared transcriptome of *ade8* during purine starvation with the one from stationary phase cells (Geodataset GSE111056, Parenteau et al., 2019). We choose relevant RNA sequencing datasets acquired using Illumina sequencing platform with at least n=2 replicates for each condition from GEO. We used the same data handling procedure as in our expression data (as described in Supplementary materials 4). When comparing both data sets, large amount of genes are simultaneously upregulated and downregulated in both conditions (see Fig 8 and full list of gene expression analyses in Supplementary materials 3). When plotted on cell map, downregulation of genes associated with translation is most obvious (Figure 8B). On another hand, upregulated genes do not show such clear clustering. Colocalization in peroxisome connected genes can be observed. We performed analyses of GO term enrichment within genes that are upregulated in purine starvation and stationary phase cells. In both cases, genes related to oxidoreductase activity and trehalose metabolism were enriched among other upregulated genes.

**Figure 8.**
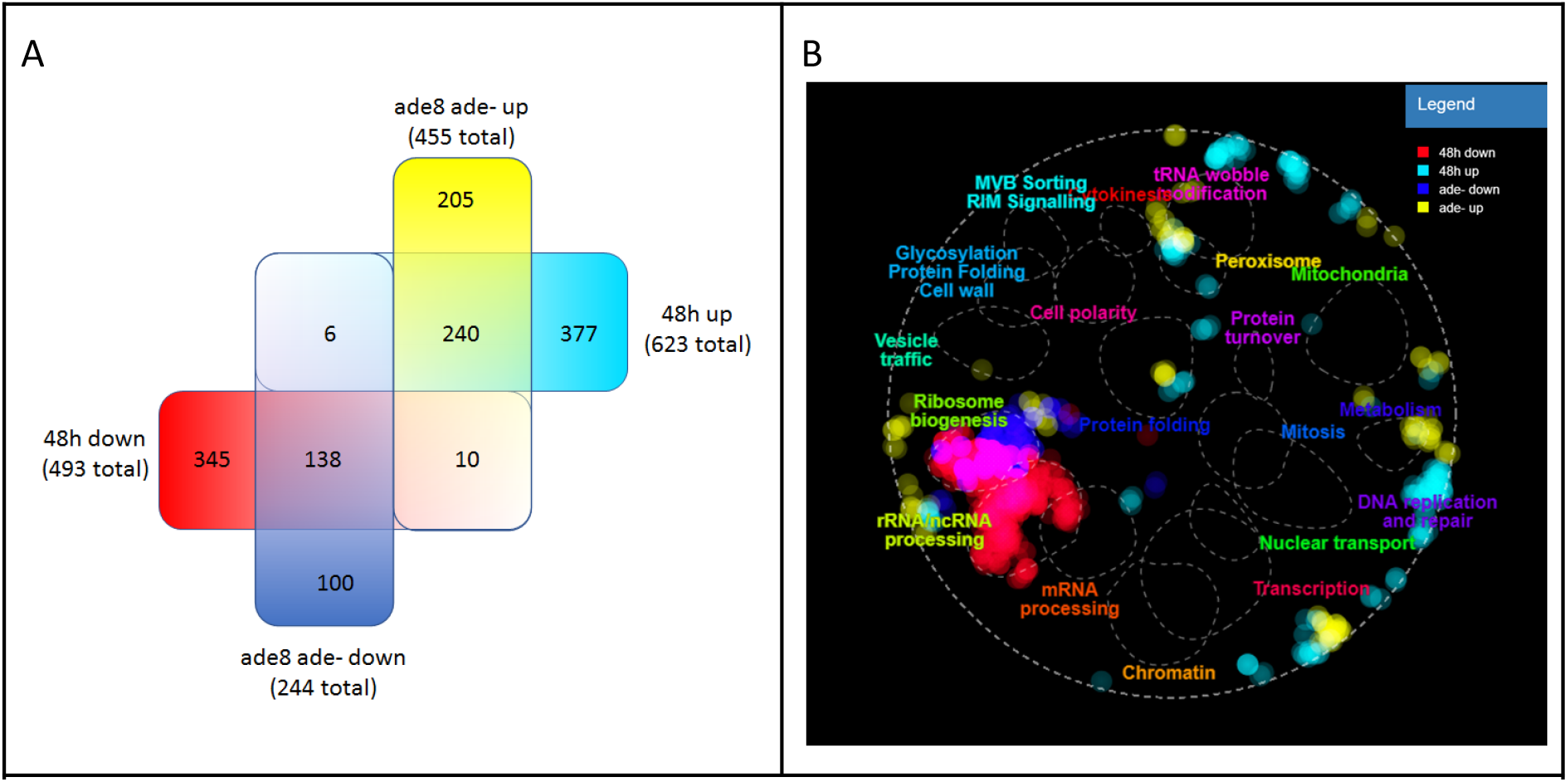
Comparison of gene expression of *ade8* strain in SDade- media after 48h and JPY10I strain (*MAT**a***/α *ura3*Δ*0/ura3*Δ*0 leu2*Δ*0/leu2*Δ*0 lys2*Δ*0/lys2*Δ*0 ADE2/ade2*Δ*::hisG HIS3/his3*Δ*200*) after 48h growth in full media (Geodata set GSE111056) **8A** Venn diagram of genes that are up regulated or downregulated with −2<logFC>2 when compared to fast growing cells. **8B** Same genes plotted with the help of TheCellMap.org

Ribosome biogenesis is an energy consuming process. In yeast rRNA represents ~60% of total transcriptional activity and ribosomal protein synthesis accounts for ~60% of PolII activity (Warner 1999). Downregulation of ribosome biogenesis is a typical process for cells in nondividing state. It is common during nutrient starvation (Conrad et al, 2014, Mayer, Grummt, 2016) and is influenced by several signalling pathways. TOR, PKA, protein C kinase and membrane secretory signalling pathways are all known to be involved in ribosome biogenesis. Nitrogen availability is mainly signalled through the TOR pathway and carbon - through PKA system. In case of amino acid starvation TOR acts via Gcn4p and Rap1p transcription factors to lower ribosome biogenesis and upregulate amino acid biosynthesis genes. (Joo et al, 2011). In case of purine starvation if analysing possible transcription factors acting, we see that targets from both - nitrogen and carbon source signalling (TOR and PKA) are affected.

## Conclusion

Budding yeast laboratory strains might contain one or many auxotrophic markers. Typically these are point or missense mutations in leucine, histidine, uracil, adenine, tryptophan synthesis pathways. Their impact on cells overall metabolism, stress resistance and transcription pattern in full media or during specific auxotrophic starvation have been studied before. Auxotrophic starvation might induce stress tolerance phenotype in the case of methionine starvation (Petti et al., 2011), but lead to stress susceptible phenotype in uracil or leucine starvation (Boer et al, 2008). Purine starvation leads to increase in desiccation, oxidative stress and acetic stress tolerance, growth of cell size, enrichment of G1/G0 cells in the culture.

We think that these distinct phenotypic changes can be explained by one or combination of following mechanisms acting during purine starvation: drop of nucleotide content in cell therefore imitates macronutrient (C, N) starvation leading to halt of the cell cycle and inducing accumulation of trehalose. Also heat shock and antioxidative systems are induced. The collection of aforementioned phenotypic markers observed during purine starvation leads to formation of strong stress resistance phenotype. In fact, we find, that constellations of phenotypic markers we observe when cells are starved for purine are the same typical for the quiescence program of *S. cerevisiae* recently reviewed by Sun and Gresham, 2021. Moreover, the quiescence program is initiated through phosphorylation of Rim15p and subsequent transcription activation of specific stress resistance genes. Previously we tested the involvement of Rim15p in development of desiccation tolerance in purine starved, *ade8* strains. We truncated C-end of the *RIM15* gene and observed that desiccation tolerance of purine starved *ade8 rim15* double mutant was lowered 100x (Ozolina et al., 2017). The similarity of purine starvation initiated phenotype to quiescence response and involvement of similar signalling pathways (Rim15p) leads us to conclude, that indeed purine starvation elicits phenotype similar if not the same as quiescence.

Our recent results demonstrate that budding yeast can adapt not only to starvation for external nutrients, like carbon or nitrogen sources, but also to deficiency of metabolic intermediates. Purine starvation is an example, where the cell compensates metabolic deficiency by “choosing” quiescence mode - stress resistance phenotype is induced to preserve the integrity and functionality of the cell. Moreover, examples from other purine auxotrophic organisms points that ability to survive purine starvation by inducing stress resistance phenotype resembling quiescence might be a universal trait of the eukaryotic cells (Kokina et al., 2019).

## Acknowledgements

Janis Liepins was supported by State Education Development agency of Republic of Latvia PostDoc Project 1.1.2/1/16/067. Agnese Kokina, Karlis Pleiko and Zane Ozolina were supported by the Latvian Council of Science, Project LZP-2018/2-0213.

