## Supplementary material 1 for "Global effects of *ade8* deletion on budding yeast metabolism": Sup1.docx

|  | CO_2_ | ethanol | acetate | glycerol | biomass | glucose uptake | Specific growth rate  μ [h^-1^] |
| --- | --- | --- | --- | --- | --- | --- | --- |
| SD+ 1 | 32,5926 | 42,5513 | 1,21479 | 1,51756 | 17,0405 | 86,5755 | 0,39875 |
| SD+ 2 | 24,8146 | 34,5129 | 1,86647 | 1,22979 | 19,2025 | 75,6468 | 0,44934 |
| SD+ 3 | 31,2419 | 31,4441 | 1,64147 | 0,92436 | 16,7734 | 81,3465 | 0,3925 |
| Ade-1 | 9,15992 | 16,2818 | 0,63756 | 2,86901 | 6,58208 | 43,0184 | 0,15402 |
| Ade-2 | 9,76494 | 16,0706 | 1,25228 | 3,71468 | 4,89126 | 40,5307 | 0,11446 |
| Ade-3 | 4,11028 | 21,9096 | 0,98627 | 4,14574 | 7,55704 | 47,6182 | 0,17683 |

Specific production (or consumption in glucose case) q values in mCmols *gDW^-1^ * h^-1^ from *ade8* cultivations in SD+ and ade- medias. Cultivations were performed in triplicates in 0.4 L reactors (starting volume 0.3 L). Samples from SD were taken after 4 hours of cultivation when max specific growth rate was reached. Samples from starving cells were harvested after 5.5 h of cultivation.
