## Supplementary material 4 for "Global effects of *ade8* deletion on budding yeast metabolism": Supplementary file 4.docx

NGS data analysis

Sequencing reads were quality filtered (Q=30), Illumina adapters and poly-A tails were removed and reads with a length of at least 100 nt were selected for further processing using cutadapt:

cutadapt -j 4 -a GATCGGAAGAGCACACGTCTGAACTCCAGTCAC -a A{100} -q 30,30 -m 100 -o out_file.format in_file.format

Reads were aligned using STAR aligner:

STAR --runThreadN 4 \

--genomeDir /c/star-genome \

--sjdbGTFfile /c/Saccharomyces_cerevisiae.R64-1-1.96.gtf \

--outFilterType BySJout \

--outFilterMultimapNmax 20 \

--alignSJoverhangMin 8 \

--alignSJDBoverhangMin 1 \

--outFilterMismatchNmax 999 \

--outFilterMismatchNoverLmax 0.04 \

--alignIntronMin 20 \

--alignIntronMax 1000000 \

--alignMatesGapMax 1000000 \

--quantMode GeneCounts \

--outSAMtype BAM SortedByCoordinate \

--limitBAMsortRAM 1287043197 \

--readFilesIn /mnt/e/liepins/liepins-ngs/53-samples-all/fg1/cut*fastq

Aligned reads were counted using HTSeq:

htseq-count -m intersection-strict -s yes -f bam -r pos Aligned.sortedByCoord.out.bam \

/mnt/h/ngs-liepins/20190424_214158/Fastq/gtf-sc/Saccharomyces_cerevisiae.R64-1-1.96.gtf \

> /c/counts.txt

edgeR was used for differential expression comparisons. Genes with less than 1 count per million (CPM) in less than 2 samples were filtered out, Benjamini and Hochberg method was used to calculate multiple comparison adjusted p-value as false discovery rate (FDR). FDR < 0.001 and logFC > 2 was set as a threshold for significance.

Workflow after adapter removal is also available on Galaxy Europe: https://usegalaxy.eu/u/karlispleiko/w/rna-seq-kp-single-read-after-cutadapt
